# Economic, social, and physiological resilience predict brain structure and cognitive performance in 9 - 10-year-old children

**DOI:** 10.1101/852988

**Authors:** Marybel Robledo Gonzalez, Clare E. Palmer, Kristina A. Uban, Terry L. Jernigan, Wesley K. Thompson, Elizabeth R. Sowell

## Abstract

While children with economic disadvantage are at risk for poorer outcomes in cognitive and brain development, less understood is the contribution of other factors in the broader socioeconomic context that may more closely index the underlying mechanisms influencing risk and resilience. We examined brain structure and cognitive test performance in association with economic disadvantage and 22 measures in the broader socioeconomic context among n = 8,158 demographically diverse 9-10-year-old children from the ABCD Study. Total cortical surface area and total cognition scores increased as a function of income-to-needs, with the steepest differences most apparent among children below and near poverty relative to their wealthier peers. We found three latent factors encompassing distinct relationships among our proximal measures, including social, economic, and physiological well-being, each associated with brain structure and cognitive performance independently of economic advantage. Our findings will inform future studies of risk and resilience in developmental outcomes for children with economic disadvantage.

## INTRODUCTION

According to the Census Bureau for 2017, 38.8% of children in the United States were living in households with economic disadvantage, ranging from deep poverty to near poverty^1^. Economic disadvantage is a socioeconomic risk factor that has been extensively reported in association with poorer outcomes in cognitive, psychosocial and physical health^2–4^. Childhood economic disadvantage has also been linked to increased risk of emergence of mental and physical health problems in adulthood^5, 6^. Most recently, studies have reported differences in characteristics of whole-brain structure among children with economic disadvantage compared to more economically advantaged peers^7, 8^. While these recent studies suggest the associations between economic disadvantage and brain structure are primarily driven by children from families with the lowest incomes, inter-individual variability among children in cognition and brain structure across the SES spectrum is not well understood^7, 8^.

Children with economic disadvantage experience more exposure to stressors that emerge from disadvantage across various economic, social, physiological, and perinatal factors that may possibly influence a child’s well-being^9–11^. For example, economic disadvantage can be accompanied by economic insecurity, such as food and housing insecurity, and adverse childhood experiences (ACEs), such as violence in the home and parental poor adaptive functioning^9, 10^. Chronic exposure to stressors can lead to a dysregulation in the stress response, influencing the activation of the hypothalamic-pituitary axis (HPA), responsible for secretion of the stress response hormones like cortisol^12^. In turn, chronic elevation of stress hormones is thought to contribute to a dysfunction in physiological systems that support healthy brain and cognitive development^2, 3, 13^.

However, economic disadvantage, as measured by family income, is a distal measure of the potential mechanisms underlying risk and resilience in developmental outcomes. Economic disadvantage is often embedded within other economic, social, physiological, and perinatal contexts that could be described by more proximal measures in the broader socioeconomic context. Importantly, while some proximal measures in the broader socioeconomic context may encompass risk for poorer developmental outcomes, other measures may index resilience in the context of economic disadvantage. For example, findings from other studies suggest social and community support, such as positive parenting and positive school environments, may be linked to resilience in developmental outcomes among children with economic disadvantage^14, 15^.

Further, adverse perinatal factors, such as low birth weight^16^ and maternal substance use^17^, have also been associated with stress dysregulation in childhood and adolescence. These same adverse perinatal factors are associated with cortical alterations^18, 19^. Children with economic disadvantage are at risk for prematurity and low birth weight ^20, 21^. Despite these connections, associations of economic, social, and perinatal risk with childhood brain and cognitive outcomes have not previously been examined within a single model^22^.

Notably, among economically advantaged and disadvantaged children, there are striking individual differences in cognitive and brain development, as well as substantial variability in the quality of economic, social, physiological, and perinatal contexts that influence children’s development. Investigating relationships among more proximal measures of the broader socioeconomic context will be crucial for understanding the contexts in which economic disadvantage is embedded and whether these contexts contribute to developmental outcomes beyond economic advantage. Here, we aimed to examine inter-individual variability across proximal measures of socioeconomic context in association with developmental measures and increase our understanding of the possible mechanisms underlying the relationship between economic disadvantage and brain and cognitive development that may ultimately be remediable with public awareness.

## RESULTS

### Greater total cortical surface area and higher total cognition scores associated with higher economic advantage

In a large sample of children 9 – 10 years of age, diverse across socioeconomic and cultural backgrounds, we tested economic disadvantage in association with developmental outcomes (see table 2 for sample demographics). Economic disadvantage was estimated using the income-to-needs percent ratio, i.e., household income relative to the federal poverty threshold for a given household size. A greater income-to-needs ratio indicated more economic advantage. Initial analyses using generalized additive mixed-effects models determined that the best fit was the smooth transformation of the income-to-needs measure, which allowed for modeling of non-linear relationships (supplementary table 2) and thus the s(income-to-needs) term was used in all models below. Effect sizes, i.e., change in R^2^, were calculated by comparing each hypothesized model to a null model that included fixed-effect covariates of age, sex, self-declared race-ethnicity, and the random effects of scanner identification number nested by family (i.e., siblings) only as predictors of each dependent variable.

**Table 1.** List of measures and groups entered into the GFA analysis.

|  |  | Measures |  |  |  |  |  |
| --- | --- | --- | --- | --- | --- | --- | --- |
| GFA Groups | Economic | Food Security | Housing Security |  | Ability to pay bills |  | Access to Medical/Dental |
|  | Parental | Parental education | Youth total caregiver acceptance |  | Youth parental monitoring |  | Dual parent households |
|  | School/Community | Youth neighborhood safety | Youth positive school environment |  | Youth school engagement |  |  |
|  | ACEs | Youth family conflict | History of traumatic event |  | Parental poor adaptive functioning |  |  |
|  | Physiological | Sufficient sleep | Body Mass Index Z-score (BMIz) |  |  |  |  |
|  | Perinatal | Total prenatal conditions | Planned pregnancy | Maternal age at birth | History of prenatal substance use | Gestational age (weeks) | Birth weight (kg) |

**Table 2.** Distributions for age, sex, and race-ethnicity by income-to-need relative to federal poverty thresholds are shown.

|  | Deep<br>Poverty<br><50% | Poverty<br>50 - <100% | Near<br>Poverty<br>100 - <200% | Mid<br>Income<br>200 - <400% | High<br>Income<br>≥ 400% | Total<br>Sample |
| --- | --- | --- | --- | --- | --- | --- |
| **Age Mean (SD)** | 9.83 (0.61) | 9.87 (0.61) | 9.92 (0.63) | 9.90 (0.63) | 9.94 (0.62) | 9.91 (0.62) |
| **Sex N (%)** |  |  |  |  |  | **N (%)** |
| Female | 303 (49.0) | 225 (46.2) | 628 (48.4) | 981 (48.0) | 1776 (47.8) | 3913 (48.0) |
| Male | 315 (51.0) | 262 (53.8) | 670 (51.6) | 1061 (52.0) | 1937 (52.1) | 4245 (52.0) |
| **Race-Ethnicity N (%)** |  |  |  |  |  |  |
| White<sup>a</sup> | 92 (14.9) | 126 (25.9) | 515 (39.7) | 1215 (59.5) | 2772 (74.7) | 4720 (57.9) |
| Hispanic | 178 (28.8) | 181 (37.2) | 362 (27.9) | 433 (21.2) | 362 (9.7) | 1516 (18.6) |
| Black<sup>a</sup> | 276 (44.7) | 133 (27.3) | 266 (20.5) | 200 (9.8) | 116 (3.1) | 991 (12.1) |
| Asian<sup>a</sup> | 4 (0.6) | 4 (0.8) | 13 (1.0) | 18 (0.9) | 91 (2.5) | 130 (1.6) |
| Other<sup>a</sup> | 68 (11.0) | 43 (8.8) | 142 (10.9) | 176 (8.6) | 372 (10.0) | 801 (9.8) |
| **Total Sample N (%)** | 618 (7.6) | 487 (6.0) | 1298 (15.9) | 2042 (25.0) | 3713 (45.5) | 8,158 (100) |
| **U.S. Population Under 18 years<sup>b</sup> %** | 8.0 | 9.5 | 21.3 | 28.9 | 32.3 | -- |

<sup>a</sup> Non-Hispanic
<sup>b</sup> U.S. Census Bureau, Current Population Survey, 2018 Annual Social and Economic Supplement.

We observed a significant non-linear relationship between income-to-needs and each developmental measure, such that both cortical surface area and cognition were more strongly related to income-to-needs among children near poverty and below, i.e., < 200% for the federal poverty level, (total cortical surface area: *F* = 34.82, *edf* = 3.77*, p* < .001, ΔR^2^_adjusted_ = 0.012, *X^2^*(1, N =8,158) = 120.66, *p* < .001; total cognition scores: *F* = 94.13, *edf* = 6.43*, p* < .001, ΔR^2^_adjusted_ = 0.064, *X^2^*(1, N =8,158) = 557.57, *p* < .001). The full model results are shown in supplementary table 4 (model 1) for total cortical surface area and in supplementary table 5 (model 1) for total cognition scores. The plot for the predicted values for the smooth term of income-to-needs for models for total cortical surface area and total cognition scores is shown in supplementary figure 1.

### Latent factors encompassing economic, social, physiological, and perinatal well-being

We implemented a Group Factor Analysis^23^ to better understand the distinct relationships among our 22 proximal measures encompassing the broader socioeconomic environment (see supplementary table 2 for the correlations among all measures). In supplementary table 3 we report the factor loading values and show consistent replication of factor loadings in two split-half samples, in a sample with singleton participants only, and in a sample randomly assigned only one participant per family. We found the first latent factor explained 13.68% of the variance across all proximal measures, and described coupled relationship among measures that indexed general social, economic, and physiological well-being (figure 2). Specifically, this latent factor loaded highly on endorsement of food security, ability to pay bills, housing security, and access to medical/dental care. Latent factor 1 also captured social well-being, loading highly on higher parental education and loading moderately on dual parent households and lower endorsement of ACEs, i.e., less endorsement across measures of parent poor adaptive functioning, history of one or more traumatic events, and family conflict. Latent factor 1 equally loaded highly on social-perinatal measures of older maternal age at birth and planned pregnancies. Lastly, latent factor 1 also encompassed physiological well-being, loading moderately on sufficient sleep, lower body-mass-index (BMI) z-scores, and on measures of physiological-perinatal health, i.e., lower prenatal conditions and lower endorsement of history of prenatal substance use. Latent factor 2 explained 6.5% of the variance across all measures and encompassed measures of youth perceived social support, loading highly on higher parental monitoring, caregiver acceptance, school engagement, and a positive school environment, and lower family conflict. Latent factor 2 also loaded to a lesser extent on lower maternal age at birth, unplanned pregnancies, and less endorsement in ability to pay bills, food and housing security (figure 2). Latent factor 3 (perinatal health) explained 5.91% of the variance and loaded on higher birth weight and gestational age, relative to lower total prenatal conditions (figure 2).

**Figure 1.**
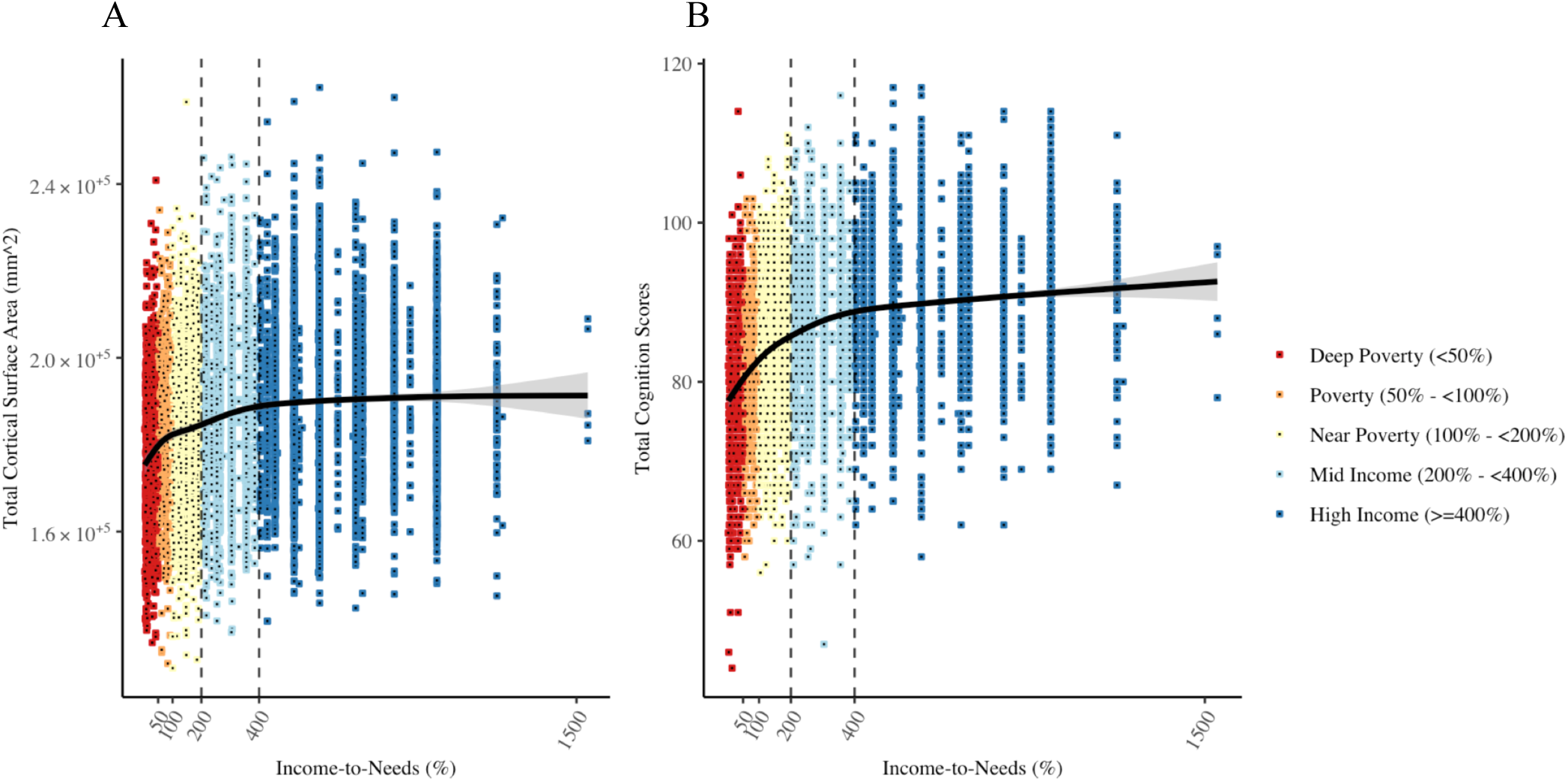
Plots showing the non-linear relationship between income-to-needs (A) total cortical surface area and (B) total cognition scores, such that increases in each developmental measure were steepest for children near poverty and below, i.e., < 200% of the federal poverty level.

**Figure 2.**
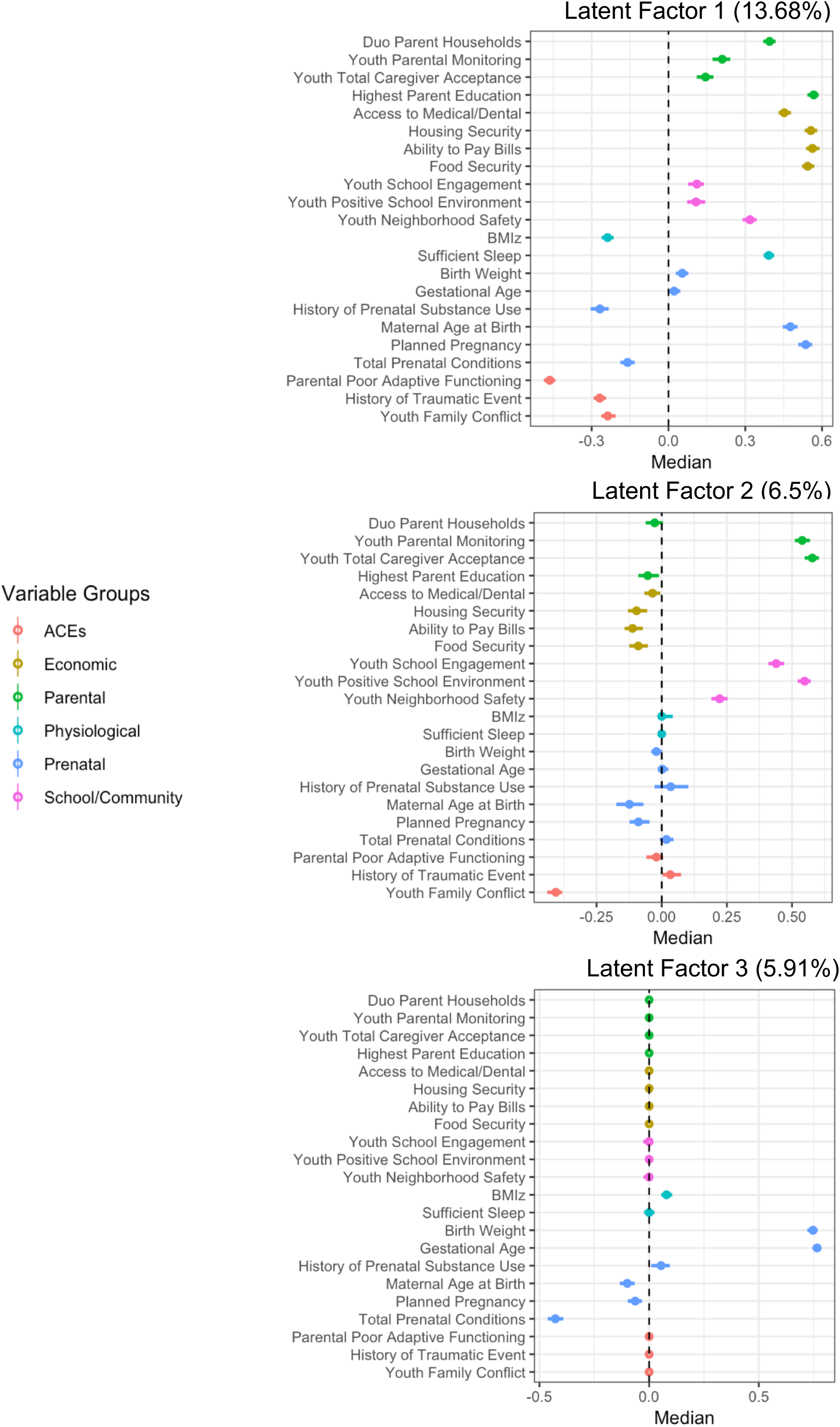
Median values for variable loadings for the three GFA latent factors shown with 95% confidence intervals, with the total variance each latent factor explained across all proximal measures.

### Latent factors positively associated with total cortical surface area and total cognition scores

The first latent factor (economic, social, and physiological well-being) was strongly associated with income-to-needs (F = 399.7, edf =7.18, p < 0.001, ΔR^2^_adjusted_ = 0.25, *X^2^*(1, N =8,158) = 2375.4, p < 0.001), suggesting that the structure of relationships among the proximal measures encompassed by this latent factor was more prevalent among families with economic advantage. The income-to-needs associations were not significant for the second latent factor of youth perceived social support (p = .97), nor for the third latent factor of perinatal well-being (p = 0.90). In separate analyses for each latent factor, adjusting for fixed covariates of age, sex, self-declared race-ethnicity, s(income-to-needs), and random effects of scanner identification number nested by family, each latent factor was positively associated with total cortical surface area (latent factor 1: economic, social, and physiological well-being, β (CI) = 0.086 (0.06, 0.112), ΔR^2^_adjusted_ = 0.014, *X^2^* (2, N =8,158) = 162.68, p < 0.001; latent factor 2: social support, β (CI) = 0.033 (0.012, 0.053), ΔR^2^_adjusted_ = 0.012, *X^2^* (2, N =8,158) = 130.46, p < 0.001; latent factor 3: perinatal health, β (CI) = 0.123 (0.101, 0.145), ΔR^2^_adjusted_ = 0.027, *X^2^* (2, N =8,158) = 236.03, p < 0.001); and each was also positively associated with the total cognition scores (latent factor 1: economic, social, and physiological well-being, β (CI) = 0.149 (0.122, 0.176), ΔR^2^_adjusted_ = 0.076, *X^2^*(2, N =8,158) = 672.81, p < 0.001; latent factor 2: social support, β (CI) = 0.049 (0.027, 0.071), ΔR^2^_adjusted_ = 0.066, *X^2^* (2, N =8,158) = 576.89, p < 0.001; latent factor 3: perinatal health, β (CI) = 0.075 (0.052, 0.098), p < 0.001, ΔR^2^_adjusted_ = 0.071, *X^2^* (2, N =8,158) = 597.95, p < 0.001). The effect sizes for models for income-to-needs and each latent factor predicting total cortical surface area and total cognition scores are plotted in figure 3. Importantly, these associations were significant when including income-to-needs in the models, which demonstrates that variability in individual differences in the developmental measures was statistically attributable to these proximal measures above and beyond economic advantage. The full model results are in supplementary tables 4 and 5.

**Figure 3.**
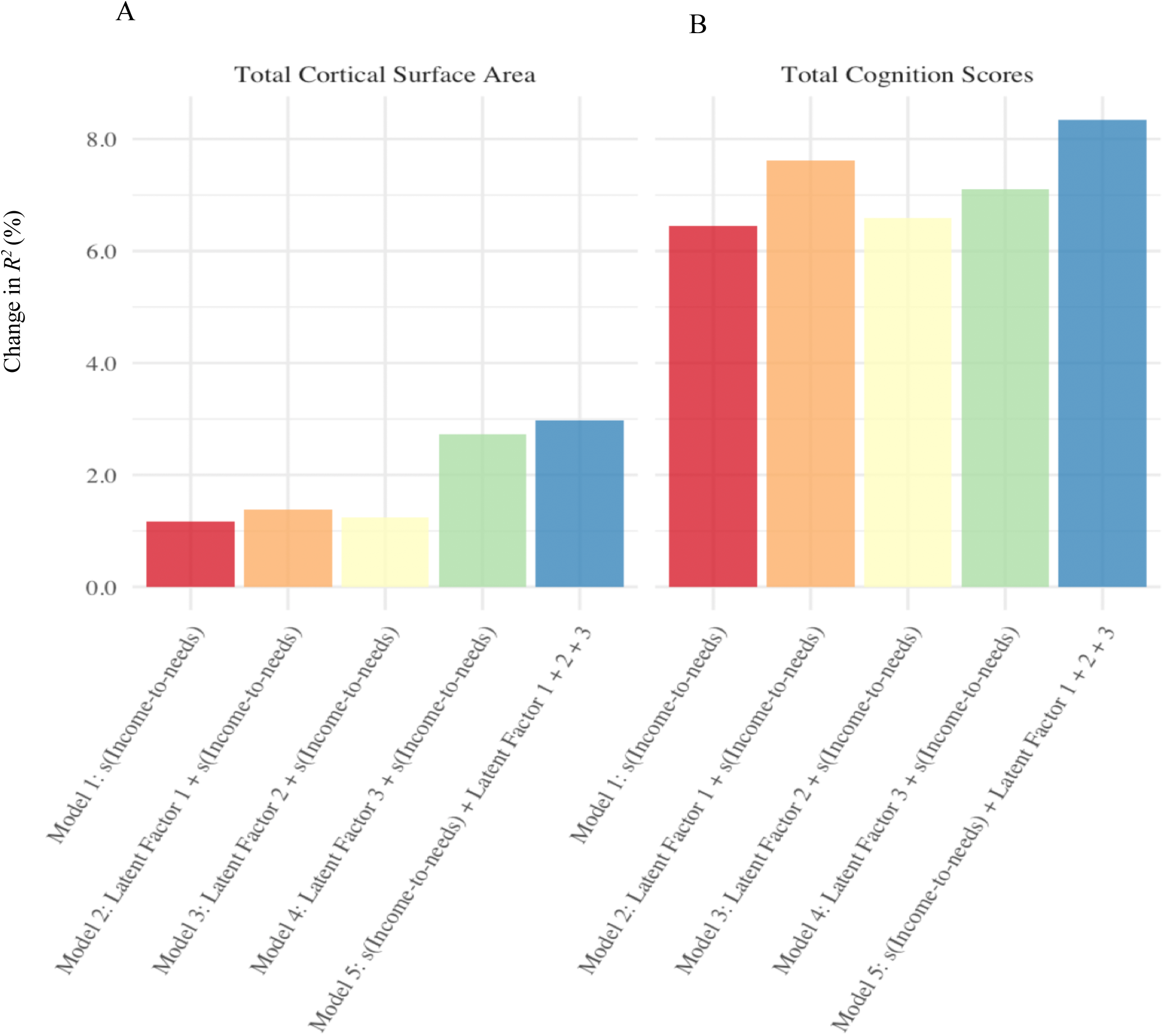
For each developmental measure (A) total cortical surface area and (B) total cognition scores, effect sizes are shown as the percent of variance statistically attributable to income-to-needs only, each latent factor, and to the additive effect of income-to-needs and all latent factors combined. Change in adjusted *R^2^* was calculated by comparing each separate model to the null model (fixed and random effects only).

### Intercation of income-to-needs and latent factor 1 on total cognition scores

To determine if there were any interactions between income-to-needs and the latent factors predicting total cortical surface area and total cognition scores, we generated a categorical variable of income-to-needs based on federal guidelines (deep poverty: <50; poverty: 50 - <100; near poverty: 100 - <200; mid income: 200 - <400; higher income: >=400). There was a significant interaction of the categorical variable of income-to-needs by latent factor 1 scores on total cognition scores (ΔR^2^_adjusted_ = 0.003, *X^2^*(4, N =8,158) = 34.8, *p* < 0.001). To interpret the interaction, we plotted the latent factor 1 scores predicting total cognition scores by income-to-needs groups (figure 4). Interestingly, the interaction plot suggess that on average, there were no apparent differences in cognition scores between income-to-needs groups among children with higher latent factor 1 scores. This suggests that children in near poverty and below with a higher relative endorsement of economic, social, and physiological well-being (as represented in latent factor 1), show comparable total cognition scores relative to their higher income peers. There was no significant interaction for latent factor 1 with income-to-needs groups on total cortical surface area (*X^2^*(4, N =8,158) = 3.76, *p* = 0.44), nor any significant interactions of latent factor 2 or latent factor 3 with income-to-needs groups on total cortical surface area (*X^2^*(4, N =8,158) <= 5.66, *ps* > 0.22) or on total cognition scores (*X^2^*(4, N =8,158) <= 4.13, *ps* > 0.39).

**Figure 4.**
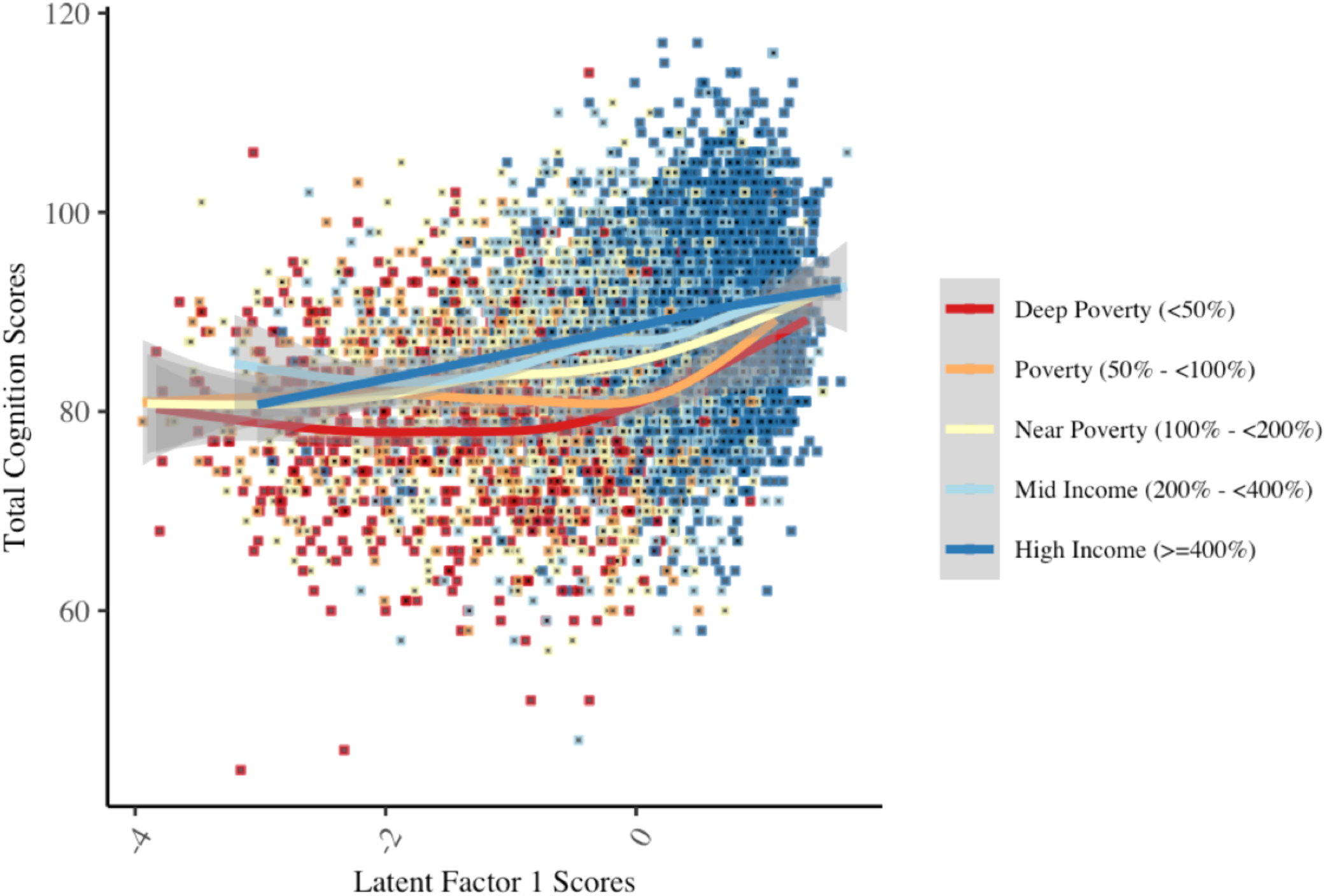
The plot of the interaction of income-to-needs by latent factor 1 scores in association with total cognition scores shows differences in total cognition scores between income-to-need groups varied as a function of latent factor 1 scores. While total cognition scores steadily increased with higher latent factor 1 scores for children in mid to high income households, total cognition scores for children in poverty (<100%) showed a protracted shift in scores, revealing an advantage in total cognition scores for children in mid to high income households for middle-range latent factor 1 scores. Importantly, the gap in total cognition scores between children in poverty relative to children in mid to high income households narrowed for children with higher latent factor 1 scores (i.e., higher endorsement of economic, social, and physiological well-being).

### Cortical Surface Area Effect Size Maps

A vertex-wise mass univariate analysis across the surface of the cortex was conducted to visualize the predictive effect of each independent variable, income-to-needs and each of the latent factors, on surface area at each vertex (figure 5). Figure 5A shows the vertex-wise association between income-to-needs (non-transformed) and surface area. Figures 5B-D show the vertex-wise association between each latent factor and surface area (in separate models) all including income-to-needs and the other latent factors as covariates. They therefore display the unique variance in surface area predicted by each latent factor independent of income-to-needs and the other orthogonal latent factors. The maximum vertex-wise beta coefficients for each predictor were β = 0.10 for income-to-needs, β =0.096 for latent factor 1, β = 0.052 for latent factor 2 and β = 0.17 for latent factor 3. The distribution of effect sizes across the cortex for income-to-needs and each of the latent factors seemed to be continuous and distributed across the cortex. The associations between surface area and latent factor 2 showed the smallest effect sizes and only a small number of vertices survived correction for multiple comparisons (see supplementary figure 4 for maps of FDR-corrected p-values). However, we cannot infer any causality or directionality from these observational associations.

**Figure 5.**
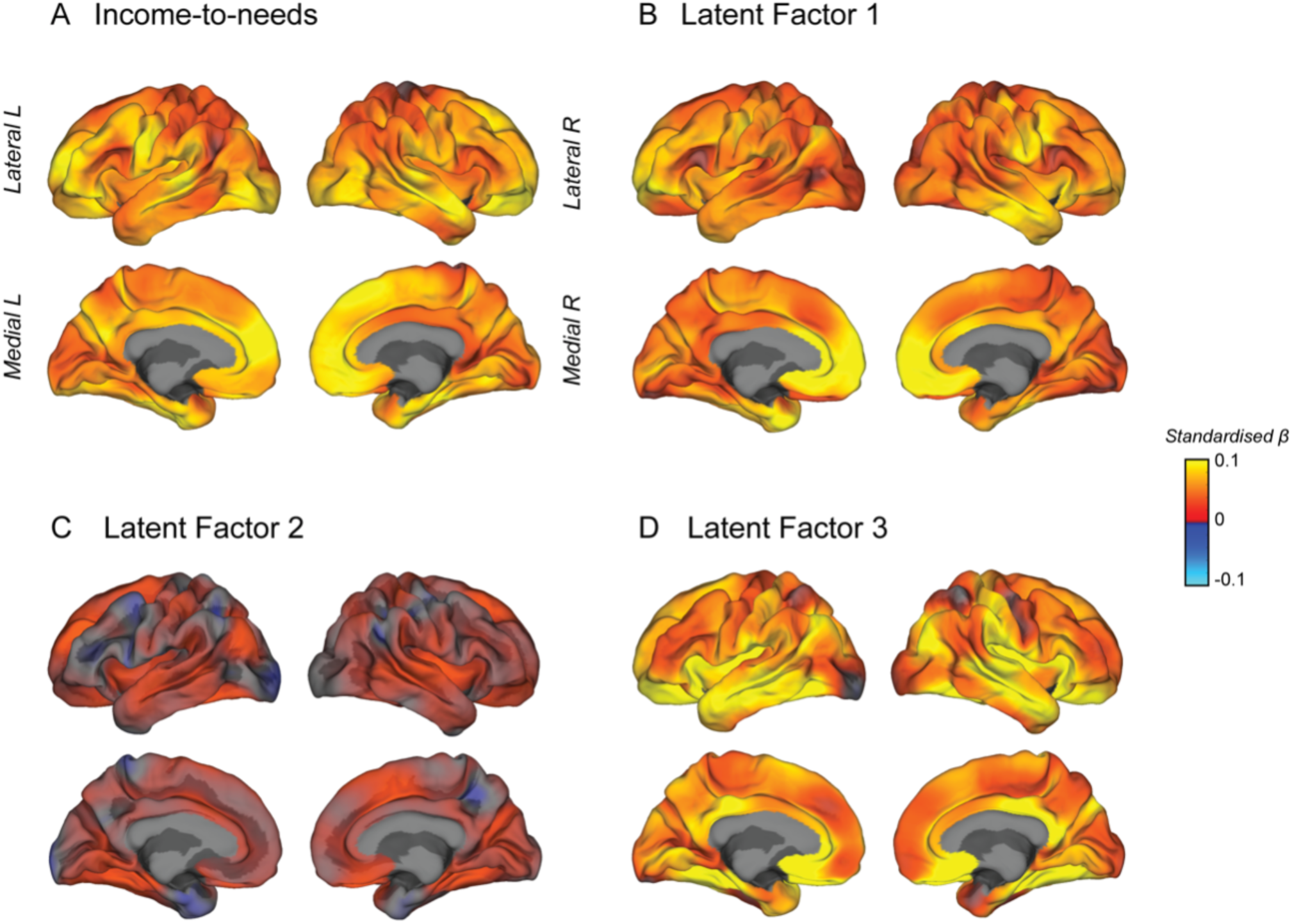
Mass univariate vertex-wise estimated effect size maps predicting surface area from each independent variable (A: income-to-needs, B: latent factor 1, C: latent factor 2 and D: latent factor 3) were created using general linear models at each vertex controlling for age, sex, race/ethnicity, and scanner. Maps B-D also included income-to-needs and the other latent factors as additional covariates such that these maps show the unique contribution of each latent factor and surface area. The maps show unthresholded standardized beta coefficients. All of the independent variables showed positive effects with surface area.

## DISCUSSION

Socioeconomic factors (e.g., family income and poverty) have long been known to impact cognitive development and school performance, with more recent research relating family income and parental education to brain development^7, 8^. Our results from a large demographically diverse cohort illuminate other factors in the broader socioeconomic context that may impart risk or resilience for negative brain and cognitive outcomes in American youth. We found associations between income-to-needs and total cortical surface area and cognitive performance, as well as independent contributions from other economic, social, physiological and perinatal measures hypothesized to be related to economic disadvantage and measures of development. We have replicated previous findings showing a non-linear relationship between income-to-needs and developmental measures, with the largest associations among children with the most economic disadvantage^7, 8^. We then examined the latent structure across 22 variables hypothesized to be associated with economic disadvantage to understand how multidimensional relationships among these more proximal measures associated with individual variability in total cortical surface area and total cognition scores beyond income-to-needs.

Each of the latent factors appeared to approximate separable relationships among our proximal measures that encompassed: (1) a general factor of economic, social and physiological well-being; (2) youth perceived social support; and (3) perinatal well-being. Each latent factor showed positive associations with each of the developmental measures. Associations between each latent factor and the brain and cognitive outcomes were significant even when including income-to-needs in the model as a covariate. Initial analyses suggested that the relative contribution of measures described by latent factor 1 (economic, social, physiological well-being) were particularly prevalent among economically advantaged children. However, we found total cognition scores varied as a function of latent factor 1 scores between income-to-needs groups. Notably, children in poverty who had higher relative endorsement of economic, social, and physiological well-being, on average, showed comparable cognition scores to more economically advantaged peers. It is likely that higher parental education affords children with more opportunities for enriching learning and recreational activities, such as participating in music or sports^24, 25^. Children in poverty with higher latent factor 1 scores, were in households with potentially more enriching opportunities (i.e., higher parental education), higher availability of parent resources (i.e., higher maternal age at birth, planned pregnancy, and dual parent households), and in secure environments that provided basic needs (i.e., food and housing security, ability to pay bills, access to medical/dental care, and sufficient sleep), as well as in healthy socio-emotional home environments (i.e., better parent adaptive functioning, no history of traumatic events, and low family conflict). While the associations between latent factor 2, youth perceived social support, and each developmental measures were moderate, our findings suggested that having a positive family and community environment was associated with positive developmental outcomes, even when coupled with some degree of risk, i.e., young maternal age at birth, unplanned pregnancies, lower endorsement of ability to pay bills and food and housing security.

While studies in children often index socioeconomic disadvantage using measures of family income and parental education, each measure studied here is thought to represent a different component of socioeconomic disadvantage by which family environments and developmental outcomes in children are influenced differently^26^. Given the limited number of studies on brain structure in children and socioeconomic disadvantage, it is not yet clear how each component may uniquely contribute to differences in brain structure^7, 8^.

Previous studies reporting on the association between family income and cortical surface area, suggested a regional specificity^7, 8^. However, those studies were with smaller sample sizes. In the present study, with increased sample size and power for detection, we see that the vertex-wise cortical surface area associations for income-to-needs and the latent factors are much more continuous and distributed across cortex, and we therefore cannot infer any regional specificity. It is unclear whether the regional distribution of effect sizes for cortical surface area associations are distinct between income-to-needs and the latent factors, which would be suggestive of differences in underlying mechanisms by which income-to-needs and the latent factors associate with total cortical surface area. However, such hypotheses can be tested in future research using multivariate analyses. Such studies could help unravel whether there are independent neurobiological mechanisms contributing to individual variability in brain structure and function arising from different socioeconomic or perinatal factors.

### Limitations

Although the composition of the study sample analyzed is overrepresented in the number of households in the higher income range relative to the population income distribution in the United States, our study sample includes a larger representation of children with economic disadvantage than previous studies^27^. The duration and extent under which children in this cohort have experienced economic and social adversity during their early childhood is not yet known. While it is challenging to differentiate between transitory poverty and chronic poverty, previous literature suggests that even children who have experienced transitory poverty have poorer outcomes compared to children who never experienced poverty^9, 28^. The relative effect sizes are small, although this is perhaps to be expected from studies examining behavioral and brain outcomes with large samples, given the heterogeneity in individual differences in the population being studied as well as the wide range of factors that could influence development. There are many other risk factors that are closely related to economic disadvantage not directly examined in this study, such as the child’s mental health and environmental toxins^3^. In addition, there are many other experiences that may contribute to resilience in developmental outcomes among children with economic disadvantage, including participation in enriching activities like art, music, and sports, that were not considered in our analysis. Future studies should examine whether there are measurable differences in the quality and access of enriching activities that stimulate learning at all levels of parental education, and whether participation in enriching activities for children among lower educated parents can be linked to positive developmental outcomes.

### Conclusion

Given the complexity of the relationships among measures of risk and resilience for children with economic disadvantage, it has been difficult to understand how such relationships between various economic, social, physiological, and perinatal factors contribute individually or multiplicatively in explaining differences in developmental outcomes^24, 29, 30^. We report findings from a comprehensive set of analyses that examined the extent to which separable relationships among measures of more proximal aspects of economic, social, physiological and perinatal well-being related to economic advantage and developmental measures in a large sample of healthy children. Our study reports timely findings that point to future areas of research to help identify factors, that if targeted appropriately with interventions, could possibly reduce risk and promote resilience for children experiencing economic disadvantage.

## Acknowledgements

The authors wish to thank the youth and families participating in the Adolescent Brain Cognitive Development (ABCD) Study and all ABCD staff. We thank Eric Kan for technical support. Data used in the preparation of this article were obtained from the Adolescent Brain Cognitive Development (ABCD) Study (https://abcdstudy.org), held in the NIMH Data Archive (NDA). This is a multisite, longitudinal study designed to recruit more than 10,000 children age 9-10 and follow them over 10 years into early adulthood. The ABCD Study is supported by the National Institutes of Health and additional federal partners under award numbers U01DA041022, U01DA041028, U01DA041048, U01DA041089, U01DA041106, U01DA041117, U01DA041120, U01DA041134, U01DA041148, U01DA041156, U01DA041174, U24DA041123, U24DA041147, U01DA041093, and U01DA041025. K.A.U. was supported by K01AA026889. A full list of supporters is available at https://abcdstudy.org/federal-partners.html. A listing of participating sites and a complete listing of the study investigators can be found at https://abcdstudy.org/Consortium_Members.pdf. ABCD consortium investigators designed and implemented the study and/or provided data but did not all necessarily participate in analysis or writing of this report. This manuscript reflects the views of the authors and may not reflect the opinions or views of the NIH or ABCD consortium investigators.

## Author Contributions

M.R.G. wrote the first draft of the manuscript. C.E.P, K.A.U., T.L.J., W.K.T., and E.R.S. edited and contributed text to the final manuscript. E.R.S., T.L.J, K.A.U., W.K.T., and C.E.P. encouraged and contributed ideas for M.R.G. to investigate and develop the theoretical framework. C.E.P. supported M.R.G in implementing the R code for the group factor analysis. C.E.P. generated the surface area effect size maps. M.R.G performed all other analyses. W.K.T. supervised all analyses. All authors discussed the results and contributed to the final manuscript.

## Methods

### Participants

Data used here were obtained from the Adolescent Brain Cognitive Development (ABCD) Study. The ABCD data repository grows and changes over time. The ABCD 2.0.1 data release was downloaded from the NIMH Data Archive ABCD Collection (10.15154/1504041) and contained baseline data for a total of N = 11,875 children ages 9 – 10 years old. Baseline data that passed quality assurance and had complete cases for FreeSurfer imaging data, demographic measures, and environment measures, were included in the analyses for a total of N = 8,158. Participants that had any incomplete data across all measures were excluded (See supplementary table 1 for details).

The recruitment strategy has been described in detail previously^31^. Children were recruited from 22 study sites and ABCD is following children at 21 study sites across the United States. A school-based recruitment strategy was developed to achieve a cohort of families that was diverse in income, race-ethnicity, and cultural background and has been described in detail by Garavan, et al., (2018). Demographic information for age, sex (female: 1, male: 0), and race/ethnicity were examined. Race/ethnicity was recoded to include 5 categories: Hispanic, and non-Hispanic White, Black, Asians, and more than one race.

### Economic advantage: Income-to-needs

Gross household income and the number of household members was reported by the study caregiver in the Parent Demographics Survey. Income was reported in categories^1^, and was adjusted to the median for each category. The income-to-needs ratio was calculated for each participant by dividing the household income median by the corresponding 2017 federal poverty level based on the Department of Human and Health Services (HHS) poverty guidelines^32^ for the reported household size. The HHS federal poverty level is the necessary income needed for a family of a given size to meet the cost of living, including shelter, food, clothing, transportation, and other necessities and determines eligibility for federal government benefit programs.

### Proximal measures in the broader socioeconomic environment

We examined 22 measures across economic, social, physiological, and perinatal well-being that we hypothesized to be related to income-to-needs and developmental measures. A complete list of measures examined is shown in table 1. We categorized the 22 measures into 6 groups thought to represent constructs that would capture within-group variability across the range of income-to-needs, specifically: (1) economic factors, (2) parental characteristics, (3) school/community environment, (4) risk for adverse childhood experiences (ACEs), (5) physiological health, and (6) perinatal well-being. Importantly, dual parent household was defined by the study caregiver report of whether he/she had a partner who was involved in at least 40% or more of the daily activities of the child. Economic security was measured by a set of questions that determined food security, housing security, ability to pay bills, and access to medical or dental care. Highest parental education was from parent report of highest education attained among both caregivers when available. For detailed information for each specific variable name and transformation, as well as missing and excluded values, see supplementary table 1. A detailed description of the ABCD baseline protocol and rationale for inclusion of measures on demographics, culture and environment have been reported previously^33, 34^.

### Cognition scores

The NIH Toolbox® cognition battery^35, 36^ was administered as part of the ABCD study baseline neurocognition battery^37^. From the NIH Toolbox® cognition battery, total cognition composite scores were examined. Total cognition composite scores are derived from seven tasks within the NIH Toolbox® cognition battery that assess working memory and categorization, information processing, flexible thinking and set shifting, visuospatial sequencing, cognitive control, reading ability and verbal intellect.

### Image acquisition and processing

The imaging procedures for ABCD imaging acquisition and preprocessing have been described previously^38^. Briefly, each site applied a standardized MRI protocol that included a T1 weighted scan. All imaging data was processed using FreeSurfer pipelines and procedures implemented by the ABCD Data Informatics and Resource Center (DAIRC)^39^. A 3D model of the cortical surface was constructed for each subject. Cortical surface area was calculated by mapping a standardized tessellation to the native space of each subject using a spherical atlas registration, which matched the cortical folding patterns across subjects. Surface area of each vertex was calculated as the area of each triangle. This generated a continuous vertex-wise measure of relative areal expansion or compression. Cortical maps were smoothed using a Gaussian kernel of 20mm full-width half maximum (FWHM) and mapped into standardized spherical atlas space. Vertexwise data for all subjects for each morphometric measurement were concatenated into matrices in MATLAB R2017a and entered into general linear models for mass univariate statistical analysis using custom written code.

### Mass univariate effect size estimation for cortical surface area

Vertexwise imaging data was obtained from the ABCD 2.0.1 fixed release and was available for 11,536 participants. Imaging data that did not pass quality assurance were excluded from our analyses using the FreeSurfer quality control variable for the ABCD baseline tabulated dataset. A total of 8,158 participants who had complete vertexwise data and complete data on all other behavioral measures were included in the vertexwise surface area analyses. To measure the vertexwise effects of income-to-needs, we conducted a GLM at every vertex predicting income-to-needs from surface area. The following fixed effects were included as covariates of no interest: age, sex, scanner identification number and race-ethnicity. To determine the vertexwise effects uniquely predicted by each latent factor from the GFA we conducted the same mass univariate vertexwise analysis including additional fixed effects of income-to-needs and the other respective latent factors. All behavioral and imaging variables were standardized with zero mean and unit variance before analysis. All estimated effect size maps show the mass univariate standardized beta coefficients. Additional maps were created showing the distribution of mass univariate p-values across the scalp adjusted for a false discovery rate (FDR) of 5% using the Benjamini-Hochberg procedure implemented in Matlab 2017a using the ‘mafdr’ function. All p-value maps were thresholded based on an alpha level of adj-p<0.05.

### Statistical analysis

All statistical analyses were done in R (3.4.4)^40^ using R studio (1.1.463)^41^. A Group Factor Analysis^23^ was implemented using the R-package GFA^42^. Generalized Additive Mixed-Effect Models (GAMMs) were fitted using the R-package gamm4^43^ to test income-to-needs and latent factor associations with total cortical surface area and cognition scores. Only participants with complete data across all 22 measures, demographic covariates, and dependent variables were included in the analyses (supplementary table 1). Continuous measures were standardized to a zero mean and unit variance. First, we tested the associations between income-to-needs and total cortical surface area and cognitive performance. Second, we conducted a group factor analysis to describe patterns of relationships among measures hypothesized to broadly encompass socioeconomic context across the entire range of income-to-needs. Third, we tested the latent factor associations with income-to-needs and each developmental measure to examine whether patterns of relationships among variables were related to income-to-needs and whether they predicted total cortical surface area and total cognition scores. Lastly, we examined interactions between income-to-needs groups and latent factors on total cortical surface area and total cognition scores.

#### Group Factor Analysis (GFA)

A GFA solution identifies linear latent factors that describe relationships among grouped variables, while also taking into account dependencies between groups. GFA is similar to a Bayesian exploratory factor analysis, except unique to the GFA approach is the implementation of a structural sparsity prior that allows modeling of the dependencies between groups. The GFA outputs linear factors that contain a projection vector comprised of the measures with non-zero loadings for that factor. All 22 measures (described in table 1) were submitted to a GFA. To test the stability and robustness of the latent factors, we completed 10 different iterations of the GFA. Robust latent factors were chosen based on latent factor loadings that met a 0.9 correlation threshold across all 10 iterations. Robust factor loadings across all 10 GFA iterations were averaged. Separate robust GFAs were examined in split-half samples to test replication of the latent factor loadings. Robust GFA latent factors accounting for more than 5% of the GFA variance were chosen.

#### Testing Associations

Null GAMMs for each dependent measure, i.e., total cortical surface area and total cognition scores, were constructed with only the covariates. The covariates included were the fixed effects of age, sex, race-ethnicity, and random effects of scanner identification number nested by family membership. The associations of income-to-needs with the dependent measures were tested by entering the smooth term of income-to-needs and the covariates as predictors of each dependent measure and comparing this model to the null model (covariates only). The associations between the latent factor variables and the dependent measures were tested by entering the latent factors, income-to-needs and covariates as predictors of each dependent measure, and these models were compared to the null model (covariates only). Each sequential model comparison was tested using a likelihood ratio test with the “anova” function in R. The variance statistically attributable by each model was interpreted as the change in *R^2^* between model comparisons and significance was determined by the chi-square (*X^2^*) statistic and corresponding *p*-value. Since all continuous measures were standardized, the model regression coefficients were interpreted as standard betas.

## Footnotes

1 Income bins: Less than $5000, $5000 to $11999, $12000 to $15999, $16000 to $24999, $25000 to $34999, $35000 to $49999, $50000 to $74999, $75000 to $99999, $100000 to $199999, $200000 and greater.

## References

1. US Census Bureau. Current Population Survey Annual Social and Economic Supplement. (2018).

2. McLoyd, V. C. Socioeconomic Disadvantage and Child Development. Am. Psychol. 53, 185–204 (1998).

3. Evans, G. W. The Environment of Childhood Poverty. Am. Psychol. 59, 77–92 (2004).

4. Farah, M. J. et al. Childhood poverty: Specific associations with neurocognitive development. (2006). doi:10.1016/j.brainres.2006.06.072

5. Jensen, S. K. G., Berens, A. E. & Nelson, C. A. Effects of poverty on interacting biological systems underlying child development. The Lancet Child and Adolescent Health 1, 225–239 (2017).

6. Melchior, M., Moffitt, T. E., Milne, B. J., Poulton, R. & Caspi, A. Why Do Children from Socioeconomically Disadvantaged Families Suffer from Poor Health When They Reach Adulthood? A Life-Course Study. Am. J. Epidemiol. 166, 966–974 (2007).

7. Hair, N. L., Hanson, J. L., Wolfe, B. L. & Pollak, S. D. Association of Child Poverty, Brain Development, and Academic Achievement. JAMA Pediatr. 169, 822 (2015).

8. Noble, K. G. et al. Family income, parental education and brain structure in children and adolescents. Nat Neurosci 18, 773–778 (2015).

9. Duncan, G. J. & Brooks-Gunn, J. Family Poverty, Welfare Reform, and Child Development. Child Dev. 71, 188–196 (2000).

10. Hussey, J. M., Chang, J. J., Kotch, J. B. & St, F. Child Maltreatment in the United States: Prevalence, Risk Factors, and Adolescent Health Consequences. Pediatrics 118, (2006).

11. Kim, K. M. et al. Associations between urinary cotinine and symptoms of attention deficit/hyperactivity disorder and autism spectrum disorder. Environ. Res. 166, 481–486 (2018).

12. Vliegenthart, J. et al. Socioeconomic status in children is associated with hair cortisol levels as a biological measure of chronic stress. Psychoneuroendocrinology 65, 9–14 (2016).

13. Kim, P., Evans, G. W., Chen, E., Miller, G. & Seeman, T. How Socioeconomic Disadvantages Get Under the Skin and into the Brain to Influence Health Development Across the Lifespan. in Handbook of Life Course Health Development 463–497 (Springer International Publishing, 2018). doi:10.1007/978-3-319-47143-3_19

14. Whittle, S. et al. Positive parenting predicts the development of adolescent brain structure: A longitudinal study. Dev. Cogn. Neurosci. 8, 7–17 (2014).

15. Benard, B. Fostering Resiliency in Kids : Protective Factors in the Family, School, and Community Protective Factors : A Research Base for the Prevention Field. (1991).

16. Papadopoulou, Z. et al. Stressful Newborn Memories: Pre-Conceptual, In Utero, and Postnatal Events. Front. Psychiatry 10, 220 (2019).

17. McLachlan, K. et al. Dysregulation of the cortisol diurnal rhythm following prenatal alcohol exposure and early life adversity. Alcohol 53, 9–18 (2016).

18. Hendrickson, T. J. et al. Two-year cortical trajectories are abnormal in children and adolescents with prenatal alcohol exposure. Dev. Cogn. Neurosci. 30, 123–133 (2018).

19. Pascoe, M. J., Melzer, T. R., Horwood, L. J., Woodward, L. J. & Darlow, B. A. Altered grey matter volume, perfusion and white matter integrity in very low birthweight adults. NeuroImage Clin. 22, 101780 (2019).

20. Malecki, C. K. & Kilpatrick Demaray, M. Social Support as a Buffer in the Relationship between Socioeconomic Status and Academic Performance. School Psychology Quarterly 21, (2006).

21. Kelly, M. M. & Li, K. Poverty, Toxic Stress, and Education in Children Born Preterm. Nurs. Res. 68, 275–284 (2019).

22. Liaw, F. & Brooks-Gunn, J. Cumulative familial risks and low-birthweight children’s cognitive and behavioral development. J. Clin. Child Psychol. 23, 360–272 (1994).

23. Klami, A., Virtanen, S., Leppaaho, E. & Kaski, S. Group Factor Analysis. IEEE Trans. Neural Networks Learn. Syst. 26, 2136–2147 (2015).

24. Guo, G. & Harris, K. M. The mechanisms mediating the effects of poverty on children’s intellectual development. Demography 37, 431–447 (2000).

25. Bradley, R. H., Corwyn, R. F., McAdoo, H. P. & García Coll, C. The home environments of children in the United States Part I: Variations by age, ethnicity, and poverty status. Child Dev. 72, 1844–1867 (2001).

26. Braveman, P. A. et al. Socioeconomic status in health research: One size does not fit all. J. Am. Med. Assoc. 294, 2879–2888 (2005).

27. Compton, W. M., Dowling, G. & Garavan, H. Ensuring the Best Use of Data: 2 The Adolescent Brain Cognitive Development Study. Press (2019).

28. Smith, M. A., Makino, S., Kvetnansky, R. & Post’, R. M. Stress and Glucocorticoids Affect the Expression of Brain-Derived Neurotrophic Factor and Neurotrophin-3 mRNAs in the Hippocampus. The Journal of Neuroscience 15, (1995).

29. Whittle, S. et al. Role of Positive Parenting in the Association Between Neighborhood Social Disadvantage and Brain Development Across Adolescence. JAMA Psychiatry 74, 824 (2017).

30. Hackman, D. & Farah, M. Socioeconomic status and the developing brain. Trends Cogn. Sci. 13, 65–73 (2009).

31. Garavan, H. et al. Recruiting the ABCD sample: Design considerations and procedures. Dev. Cogn. Neurosci. 32, 16–22 (2018).

32. Department of Humuan Health Services (HSS). Poverty Guidelines | ASPE. Available at: https://aspe.hhs.gov/2017-poverty-guidelines. (2017) (Accessed: 6th November 2019)

33. Barch, D. M. et al. Demographic, physical and mental health assessments in the adolescent brain and cognitive development study: Rationale and description. Dev. Cogn. Neurosci. 32, 55–66 (2018).

34. Zucker, R. A. et al. Assessment of culture and environment in the Adolescent Brain and Cognitive Development Study: Rationale, description of measures, and early data. Dev. Cogn. Neurosci. 32, 107–120 (2018).

35. Bleck, T. P., Nowinski, C. J., Gershon, R. & Koroshetz, W. J. What is the NIH Toolbox, and what will it mean to neurology? (2013).

36. Gershon, R. C. et al. IV. NIH Toolbox Cognition Battery (CB): Measuring language (vocabulary comprehension and reading decoding). Monogr. Soc. Res. Child Dev. 78, 49–69 (2013).

37. Luciana, M. et al. Adolescent neurocognitive development and impacts of substance use: Overview of the adolescent brain cognitive development (ABCD) baseline neurocognition battery. Dev. Cogn. Neurosci. 32, 67–79 (2018).

38. Casey, B. J. et al. The Adolescent Brain Cognitive Development (ABCD) study: Imaging acquisition across 21 sites. Developmental Cognitive Neuroscience 32, 43–54 (2018).

39. Hagler, D. J. et al. Image processing and analysis methods for the Adolescent Brain Cognitive Development Study. Neuroimage 116091 (2019). doi:10.1016/j.neuroimage.2019.116091

40. Team, R. C. R: A language and environment for statistical computing. (2013).

41. Team R 2016. RStudio: Integrated Development Environment for R. RStudio,. (RStudio, Inc.).

42. Package ‘GFA’ Title Group Factor Analysis. (2017). doi:10.1109/TNNLS.2014.2376974

43. Wood, S. N. Generalized additive models: An introduction with R, second edition. Generalized Additive Models: An Introduction with R, Second Edition (Chapman and Hall/CRC, 2017). doi:10.1201/9781315370279

