## Supplementary Material for "Economic, social, and physiological resilience predict brain structure and cognitive performance in 9 - 10-year-old children"

**A** Total Cortical Surface Area Model **B** Total Cognition Scores Model

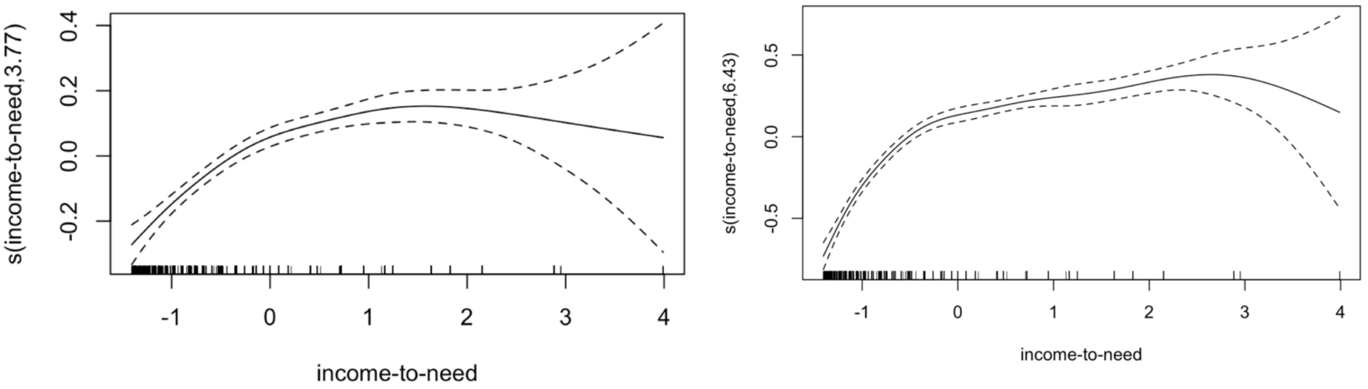

**Supplementary Figure 1.** Plots showing the predicted values for the smooth (s) term of income-to-needs for the model predicting (A) total cortical surface area (estimated degrees of freedom = 3.77) and for the model predicting (B) total cognition scores (estimated degrees of freedom = 6.43). All models controlled for fixed effects of age, sex, race/ethnicity, scanner, and random effect of family.

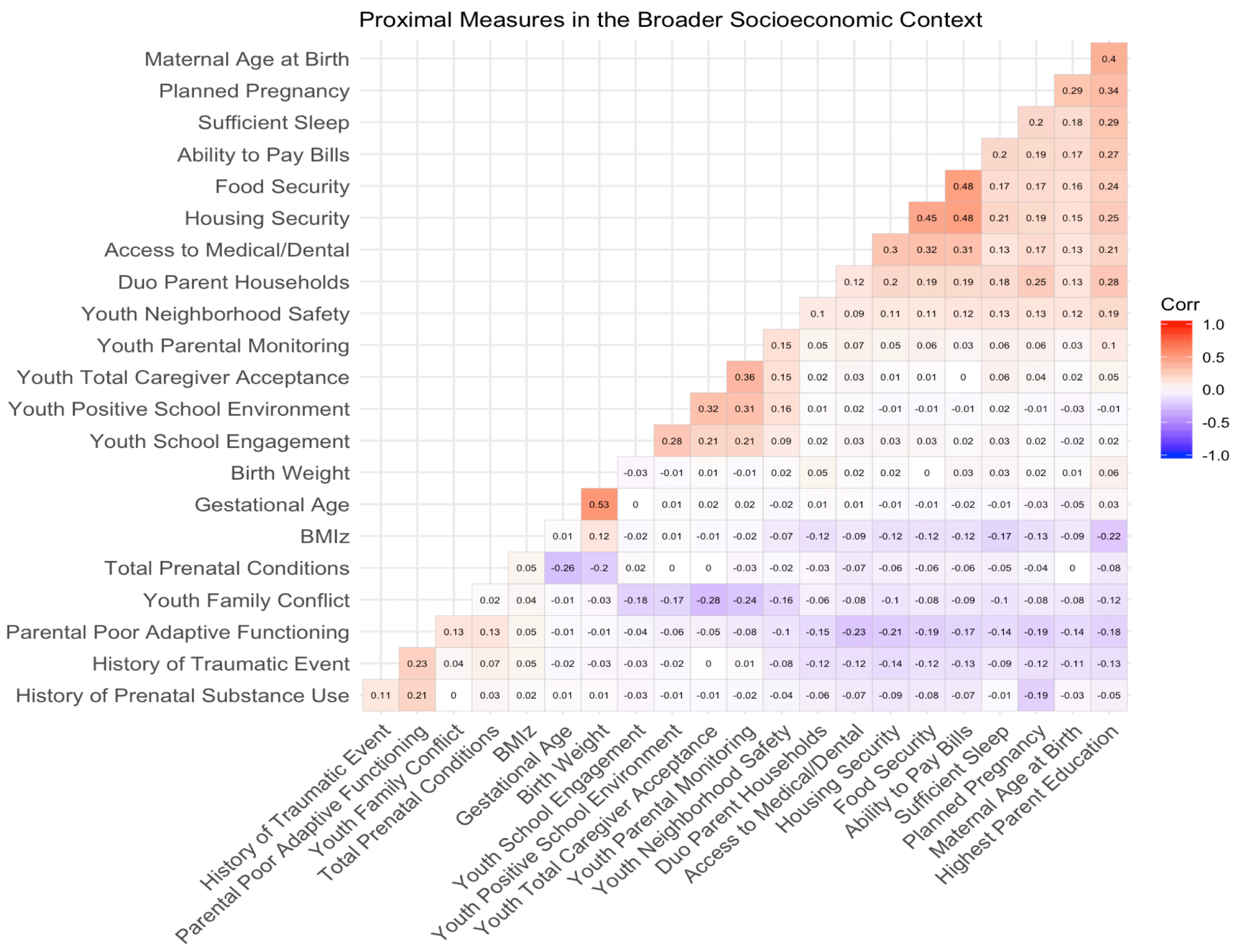

**Supplementary Figure 2.** Spearman correlation matrix showing the correlation structure for all 22 measures considered as proximal measures of the broader socioeconomic context.

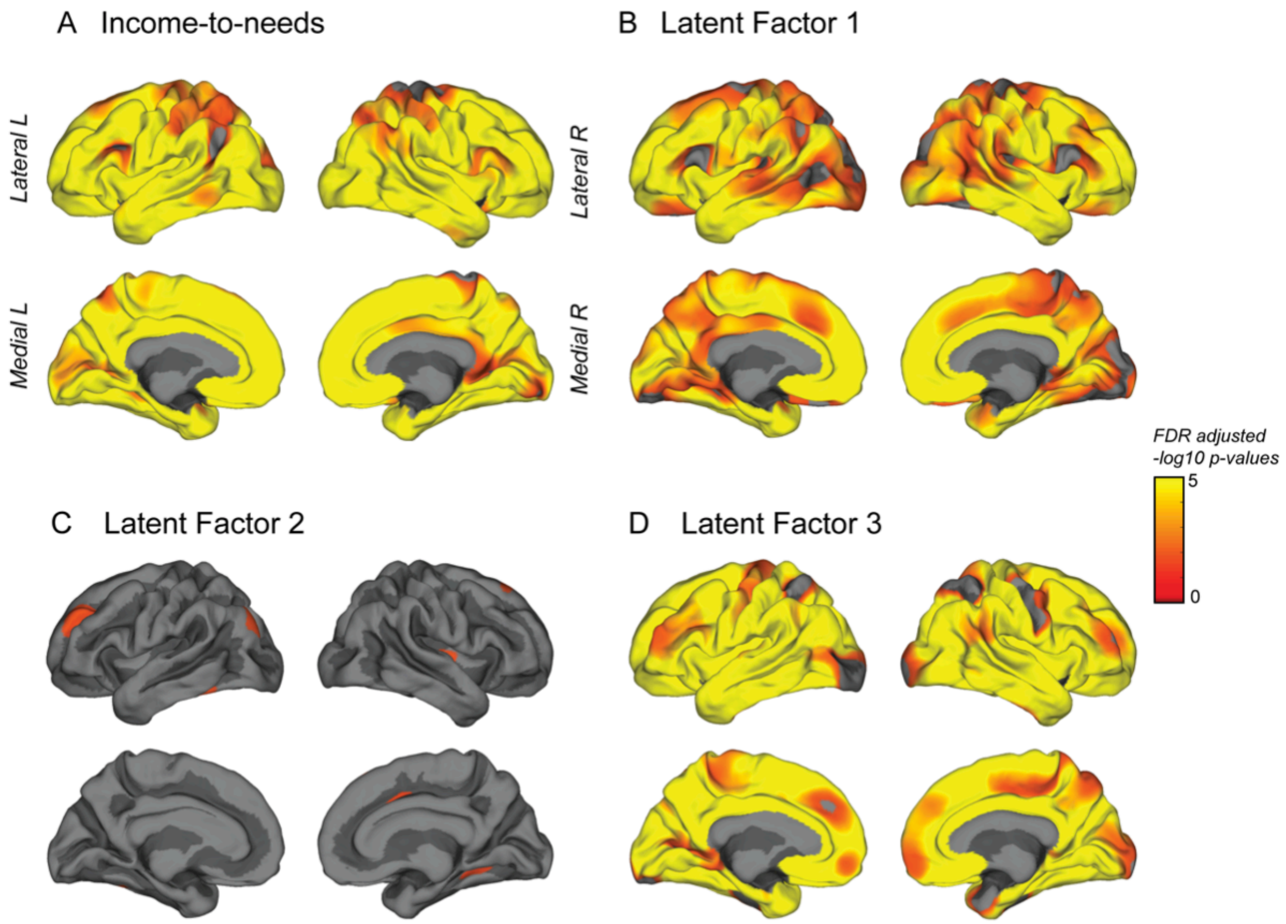

**Supplementary Figure 3.** Mass univariate thresholded vertexwise p-values, adjusted for a false discovery rate (FDR) of 5%, predicting surface area from each independent variable (A: income-to-needs, B: latent factor 1, C: latent factor 2, D: latent factor 3 at each vertex, controlling for age, sex, race/ethnicity, and scanner. Maps B-D included income-to-needs and the other latent factors as additional covariates such that these maps show the unique contribution of each latent factor and surface area.

**Supplementary Table 1.** List of all measures with corresponding NDA variable names from the ABCD 2.0 dataset.

| Group | Measure | ABCD Questionnaire | NDA variables and transformation | N Missing | N Excluded |
|---|---|---|---|---|---|
| **Economic Advantage** | | Demographic Questionnaire | nda18$demo_comb_income_v2b<br>income-recoded:<br>2500 = Less than $5,000<br>8500 = $5,000 through $11,999<br>14000 = $12,000 through $15,999<br>20500 = $16,000 through $24,999<br>30000 = $25,000 through $34,999<br>42500 = $35,000 through $49,999<br>62500 = $50,000 through $74,999<br>87500 = $75,000 through $99,999<br>150000 = $100,000 through $199,999<br>250000 = $200,000 and greater<br>NA = Refuse to answer<br>NA = Don't know | 1,018 | |
| | | 2017 Poverty Guidelines | Federal poverty guidelines [https://aspe.hhs.gov/2017-poverty-guidelines](https://aspe.hhs.gov/2017-poverty-guidelines) | -- | -- |
| | Household size | Demographic Questionnaire | nda18$demo_roster_p | 281 | 33 (values = 0 or > 20) |
| | FPL | -- | Each participant assigned Federal poverty level (FPL) based on household size and 2017 Poverty Guidelines | 281 | 33 |
| | Income-to-need | -- | (Income-recoded/FPL)*100 | 1213 | |
| **Proximal Measures of** | Food insecurity | Demographic Questionnaire | Recoded: No/Yes (0/1)<br>NA = Don't know<br>demo_fam_exp1_v2b | 77 | |

| | | | | |
|---|---|---|---|---|
| **Economic Security** | Ability to pay bills | Demographic Questionnaire | Recoded: No/Yes (0/1)<br>NA = Don’t know<br>demo_fam_exp2_v2b or demo_fam_exp5_v2b | 48 |
| | Housing Security | Demographic Questionnaire | Recoded: No/Yes (0/1)<br>NA = Don’t know<br>demo_fam_exp3_v2b or demo_fam_exp4_v2b | 63 |
| | Lack of Access to Medical/ Dental | Demographic Questionnaire | Recoded: No/Yes (0/1)<br>NA = Don’t know<br>demo_fam_exp6_v2b or demo_fam_exp7_v2b | 52 |
| **Parental** | Parental Education | Demographic Questionnaire | high.educ (DEAP)<br>Responses were re-coded to numerical values corresponding to 1 (<High School); 2 (High School); 3 (Some college); 4 (B.A.); 4 (Post Graduate) | 17 |
| | Youth Total Caregiver Acceptance | Children's Report of Parental Behavioral Inventory | crpbi_ss_studycaregiver | 36 |
| | Youth Parental Monitoring | Parental Monitoring Survey | Recoded:<br>1 = Strongly Disagree<br>2 = Disagree<br>3 = Neutral<br>4 = Agree<br>5 = Strongly Agree<br>NA = Don’t know<br>parental_monitoring_q1+<br>parental_monitoring_q2+<br>parental_monitoring_q3+<br>parental_monitoring_q4+<br>parental_monitoring_q5 | 25 |
| | Duo Parent Households | Demographic Questionnaire | demo_prnt_prtnr_v2b | 126 |
| **School/ Community** | Youth Neighborhood Safety | Youth Neighborhood Safety/Crime Survey Modified from PhenX (NSC) | neighb_phenx | 26 |

| | | | | |
|---|---|---|---|---|
| | Youth Positive School Environment | School Risk and Protective Factors Survey | Recoded:<br>1 = No!<br>2 = no<br>3 = yes<br>4 = Yes!<br>NA = Don't know<br><br>school_risk_phenx_2+<br>school_risk_phenx_3+<br>school_risk_phenx_4+<br>school_risk_phenx_5+<br>school_risk_phenx_6+<br>school_risk_phenx_7 | 28 |
| | Youth School Engagement | School Risk and Protective Factors Survey | Recoded:<br>4 = No!<br>3 = no<br>2 = yes<br>1 = Yes!<br>NA = Don't know<br>school_risk_phenx_15+<br>school_risk_phenx_17 | 27 |
| **ACEs** | Youth Family Conflict | Youth Family Environment Scale-Family Conflict Subscale Modified from PhenX (FES) | fes_y_ss_fc | 26 |
| | History of Traumatic event | Parent Diagnostic Interview for DSM-5 (KSADS) Traumatic Events | Recoded: No/Yes (0/1)<br>NA = Don't know<br>ksads_21_134_p | 185 |
| | Parental Poor Adaptive Functioning | Family History Questionnaire & Adult Self Report (total problems score) | Recoded: No/Yes (0/1)<br>(z-score(asr_scr_totprob_t)<br>+zscore(famhx_q9a_trouble_p or famhx_q9d_trouble _p<br>))/2 | 108 |
| **Perinatal** | Total Prenatal Conditions | Developmental History Questionnaire | Recoded: No/Yes (0/1)<br>NA = Don't know<br>devhx_10b_heavy_bleeding_p +<br>devhx_10c_eclampsia_p + | 712 |

| | | | | | |
|---|---|---|---|---|---|
| | | | devhx_10d_gall_bladder_p + devhx_10e_persist_proteinuria_p + devhx_10f_rubella_p + devhx_10g_severe_anemia_p + devhx_10h_urinary_infections_p + devhx_10i_diabetes_p + devhx_10j_high_blood_press_p + devhx_10k_problems_placenta_p + devhx_10l_accident_injury_p + devhx_10m_other_p | | |
| | Planned Pregnancy | Developmental History Questionnaire | Recoded: No/Yes (0/1)<br>NA = Don't know<br>devhx_6_pregnancy_planned_p | 267 | |
| | Maternal Age at Birth | Developmental History Questionnaire | devhx_3_age_at_birth_mother_p | 264 | |
| | History of Prenatal Substance Use | Developmental History Questionnaire | Recoded: No/Yes (0/1)<br>NA = Don't know<br>(devhx_8_tobacco_p or devhx_8_alcohol_p or devhx_8_marijuana_p or devhx_8_coc_crack_p or devhx_8_her_morph_p or devhx_8_oxycont_p or devhx_8_other_drugs) + (devhx_9_tobacco_p or devhx_9_alcohol_p or devhx_9_marijuana_p or devhx_9_coc_crack_p or devhx_9_her_morph_p or devhx_9_oxycont_p or devhx_9_other_drugs) | 730 | |
| | Gestational Age | Developmental History Questionnaire | 40 - devhx_12_weeks_premature_p | 173 | |
| | Birth Weight (kg) | Developmental History Questionnaire | (devhx_2_birth_wt_lbs_p + devhx_2b_birth_wt_oz_p) * 0.453592 | 525 | 2 (> 4.98 kg @ 35 weeks gestation) |

| | | | | | |
|---|---|---|---|---|---|
| **Physiological** | Sufficient Sleep | Youth Anthropometrics Modified From PhenX (ANT) survey | sleep_1_p<br>numerically recoded: 1 = less than 5 hours, 2 = 5-7 hours, 3 = 7-8 hours, 4 = 8-9 hours, 5 = 9 -11 hours | 6 | |
| | BMIz | Youth Anthropometrics Modified From PhenX (ANT) survey & SAS Program for the 2000 CDC Growth Charts[1] | anthro_height_calc *2.54 (cm)<br>anthro_weight_calc *0.453592 (kg)<br>age<br>sex | 15 | 46 (BMIz < -4) |
| **Imaging** | Total Cortical Surface Area | sMRI Part 1 | smri_area_cort.desikan_total | 341 | 462 |
| | Quality control for Freesurfer data | FreeSurfer QC | fsqc_qc | 337 | -- |
| **Cognition** | NIH Toolbox Total Computed Score Uncorrected | Youth NIH TB Summary Scores | nihtbx_fluidcomp_uncorrected | 249 | |
| **Covariates** | MRI Serial number | MRI Info | mri_info_device.serial.number | 123 | -- |
| | Race/ethnicity | DEAP | race.4level<br>demo_ethn_p<br><br>Recoded:<br>Hispanic (demo_ethn_p = 1)<br>White ((demo_ethn_p != 1 & race.4level = “White”)<br>Black ((demo_ethn_p != 1 & race.4level = “Black”)<br>Asian ((demo_ethn_p != 1 & race.4level = “ Asian ”) | 54 | |

[1] Centers for Disease and Control and Prevention. A SAS Program for the 2000 CDC Growth Charts. Available at: https://www.cdc.gov/nccdphp/dnpao/growthcharts/resources/sas.htm. (Accessed: 19th August 2019)

| | | | | | |
|---|---|---|---|---|---|
| | | | Other/Mixed ((demo_ethn_p != 1 & race.4level = “ Other/Mixed ”) | | |
| | Age | DEAP | age | 0 | -- |
| | Sex | DEAP | sex<br>Recoded:<br>F (1)<br>M (0) | 4 | -- |
| | Family | DEAP | rel_family_id | 0 | -- |

**Supplementary Table 2.** Results of the log-likelihood tests comparing the untransformed income-to-needs (Model 1) as a predictor, and the log of income-to-needs (Model 2) as a predictor, to the null model with fixed and random effects only as predictors (age + sex + race.ethnicity, random = 1| scanner/family). The models with the log of income-to-needs had the lowest AIC values suggesting these models were the best fitting models.

| | **Null Model** | **Model 1 income-to-needs (untransformed)** | **Model 2 *s*(income-to-needs)** |
|---|---|---|---|
| **Total Cortical Surface Area** | | | |
| AIC | 19378 | 19286 | 19262 |
| Chi-square | -- | 94.57 | 120.66 |
| p-value | -- | < 0.001 | <0.001 |
| **Total Cognition Scores** | | | |
| AIC | 20750 | 20360 | 20197 |
| Chi-square | -- | 392.35 | 557.57 |
| p-value | -- | < 0.001 | <0.001 |

**Supplementary Table 3.** Median latent factor loadings (95% CI) show consistent replication in latent factors across the reported robust GFA with the full sample, the robust GFAs for each split-half sample, the robust GFA with a sample that included singletons only, and the robust GFA with a sample randomly assigned only one participant per family. All robust GFAs were averaged across 10 GFA iterations.

| | **Measure** | **Full Sample N = 8158** | **Split-half sample 1 N = 3305** | **Split-half sample 2 N = 4079** | **Singletons Only N = 4079** | **Random Sample for 1 participant per family N = 6879** |
|---|---|---|---|---|---|---|
| **Latent Factor 1** | Total Prenatal Conditions | -0.16 (-0.188, -0.1315) | -0.148 (-0.188, -0.108) | -0.164 (-0.2015, -0.126) | -0.156 (-0.1845, -0.1195) | -0.16 (-0.189, -0.133) |
| | Planned Pregnancy | 0.536 (0.507, 0.5615) | 0.509 (0.468, 0.5525) | 0.552 (0.5145, 0.5835) | 0.545 (0.515, 0.574) | 0.538 (0.51, 0.568) |
| | Maternal Age at Birth | 0.476 (0.4465, 0.5045) | 0.472 (0.44, 0.5115) | 0.477 (0.4295, 0.5175) | 0.474 (0.4425, 0.503) | 0.478 (0.44, 0.51) |
| | History of Prenatal Substance Use | -0.268 (-0.304, -0.2335) | -0.255 (-0.3075, -0.203) | -0.247 (-0.3075, -0.2045) | -0.266 (-0.305, -0.224) | -0.258 (-0.298, -0.215) |
| | Gestational Age | 0.022 (-0.002, 0.046) | 0.026 (-0.0115, 0.0615) | 0.019 (-0.012, 0.053) | 0.02 (-0.011, 0.0455) | 0.024 (-0.004, 0.051) |
| | Birth Weight | 0.054 (0.029, 0.078) | 0.066 (0.03, 0.102) | 0.04 (0.007, 0.0755) | 0.054 (0.0255, 0.081) | 0.05 (0.025, 0.08) |
| | Youth Family Conflict | -0.238 (-0.2625, -0.2065) | -0.204 (-0.239, -0.155) | -0.25 (-0.294, -0.21) | -0.233 (-0.2645, -0.1985) | -0.234 (-0.269, -0.206) |
| | History of Traumatic Event | -0.268 (-0.2925, -0.2435) | -0.24 (-0.2755, -0.205) | -0.276 (-0.312, -0.2425) | -0.268 (-0.297, -0.2415) | -0.264 (-0.292, -0.238) |
| | Parental Poor Adaptive Functioning | -0.464 (-0.487, -0.442) | -0.45 (-0.484, -0.4165) | -0.454 (-0.4875, -0.42) | -0.463 (-0.487, -0.4355) | -0.458 (-0.482, -0.43) |
| | Highest Parent Education | 0.567 (0.5425, 0.5875) | 0.571 (0.5375, 0.6065) | 0.57 (0.539, 0.607) | 0.56 (0.535, 0.587) | 0.562 (0.539, 0.589) |
| | Youth Total Caregiver Acceptance | 0.144 (0.1105, 0.176) | 0.14 (0.083, 0.1865) | 0.126 (0.074, 0.173) | 0.145 (0.104, 0.1845) | 0.142 (0.109, 0.185) |
| | Youth Parental Monitoring | 0.21 (0.172, 0.2415) | 0.192 (0.142, 0.2375) | 0.202 (0.1555, 0.253) | 0.203 (0.1645, 0.244) | 0.208 (0.173, 0.245) |
| | Duo Parent Households | 0.395 (0.37, 0.4185) | 0.378 (0.3415, 0.413) | 0.421 (0.3875, 0.456) | 0.395 (0.368, 0.422) | 0.397 (0.371, 0.421) |
| | Sufficient Sleep | 0.392 (0.3715, 0.414) | 0.428 (0.391, 0.456) | 0.376 (0.3465, 0.4075) | 0.4 (0.373, 0.4255) | 0.392 (0.368, 0.418) |
| | BMIz | -0.238 (-0.2615, -0.214) | -0.254 (-0.288, -0.2205) | -0.237 (-0.2705, -0.208) | -0.241 (-0.272, -0.2115) | -0.25 (-0.28, -0.227) |
| | Youth Neighborhood Safety | 0.318 (0.29, 0.344) | 0.281 (0.252, 0.319) | 0.312 (0.2805, 0.346) | 0.32 (0.2905, 0.3485) | 0.311 (0.281, 0.34) |
| | Youth Positive School Environment | 0.108 (0.073, 0.1435) | 0.078 (0.03, 0.121) | 0.108 (0.0615, 0.1525) | 0.101 (0.0625, 0.1395) | 0.102 (0.066, 0.148) |

| | | | | | | |
|---|---|---|---|---|---|---|
| | Youth School Engagement | 0.111 (0.077, 0.139) | 0.076 (0.0365, 0.119) | 0.14 (0.098, 0.184) | 0.106 (0.073, 0.141) | 0.106 (0.073, 0.146) |
| | Food Security | 0.544 (0.522, 0.57) | 0.569 (0.5315, 0.6085) | 0.544 (0.5065, 0.581) | 0.546 (0.5215, 0.576) | 0.544 (0.522, 0.575) |
| | Ability to Pay Bills | 0.562 (0.538, 0.59) | 0.6 (0.562, 0.6355) | 0.555 (0.516, 0.588) | 0.571 (0.5455, 0.599) | 0.562 (0.536, 0.592) |
| | Housing Security | 0.556 (0.532, 0.5815) | 0.564 (0.5275, 0.596) | 0.569 (0.531, 0.605) | 0.56 (0.531, 0.5865) | 0.55 (0.521, 0.58) |
| | Access to Medical/Dental | 0.452 (0.43, 0.479) | 0.462 (0.4235, 0.495) | 0.45 (0.4135, 0.487) | 0.448 (0.42, 0.4745) | 0.448 (0.422, 0.471) |
| **Latent Factor 2** | Total Prenatal Conditions | 0.018 (-0.0075, 0.0455) | 0 (0.0015, -0.0035) | 0.012 (-0.026, 0.0485) | 0.023 (0.06, -0.011) | 0.016 (-0.005, 0.043) |
| | Planned Pregnancy | -0.09 (-0.1235, -0.0465) | 0 (0, -0.0315) | -0.086 (-0.125, -0.0405) | -0.104 (-0.0595, -0.1545) | -0.074 (-0.129, -0.043) |
| | Maternal Age at Birth | -0.124 (-0.1745, -0.07) | 0 (0, -0.0485) | -0.138 (-0.192, -0.086) | -0.124 (-0.068, -0.1835) | -0.109 (-0.17, -0.047) |
| | History of Prenatal Substance Use | 0.034 (-0.0275, 0.102) | 0 (0.0085, 0) | 0.019 (-0.0675, 0.0885) | 0.06 (0.1285, -0.024) | 0.036 (-0.005, 0.111) |
| | Gestational Age | 0.002 (-0.015, 0.0255) | 0 (0, -0.007) | 0.015 (-0.013, 0.0485) | 0 (0.0255, -0.0245) | 0.004 (-0.016, 0.03) |
| | Birth Weight | -0.02 (-0.04, 0.002) | 0 (0, -0.012) | -0.02 (-0.047, 0.01) | -0.02 (0.0025, -0.0495) | -0.011 (-0.033, 0.012) |
| | Youth Family Conflict | -0.406 (-0.4395, -0.382) | -0.422 (-0.385, -0.4665) | -0.398 (-0.4365, -0.3525) | -0.409 (-0.3735, -0.4405) | -0.406 (-0.438, -0.378) |
| | History of Traumatic Event | 0.033 (0.003, 0.0745) | 0.012 (0.0535, -0.025) | 0.048 (-0.001, 0.0935) | 0.046 (0.085, 0.006) | 0.041 (0.01, 0.076) |
| | Parental Poor Adaptive Functioning | -0.02 (-0.059, 0.0055) | -0.016 (0.025, -0.052) | -0.046 (-0.101, -0.0105) | -0.006 (0.035, -0.0475) | -0.02 (-0.054, 0.013) |
| | Highest Parent Education | -0.054 (-0.09, -0.0105) | 0.016 (0.0515, -0.018) | -0.081 (-0.129, -0.0285) | -0.037 (-0.0055, -0.072) | -0.041 (-0.074, -0.001) |
| | Youth Total Caregiver Acceptance | 0.578 (0.5485, 0.6035) | 0.597 (0.631, 0.5585) | 0.583 (0.5495, 0.624) | 0.574 (0.6035, 0.5435) | 0.591 (0.563, 0.62) |
| | Youth Parental Monitoring | 0.539 (0.5105, 0.569) | 0.566 (0.602, 0.5275) | 0.536 (0.496, 0.5705) | 0.538 (0.57, 0.5015) | 0.552 (0.521, 0.584) |
| | Duo Parent Households | -0.027 (-0.0615, 0.0045) | 0.012 (0.0615, -0.0445) | -0.047 (-0.095, 0) | -0.019 (0.017, -0.058) | -0.022 (-0.054, 0.016) |
| | Sufficient Sleep | 0 (-0.0105, 0.0025) | 0 (0.0495, 0) | 0 (-0.044, 0) | 0 (5e-04, -0.014) | 0 (-0.01, 0.004) |
| | BMIz | 0 (0, 0.0425) | 0 (0.043, -0.0015) | 0 (0, 0.0595) | 0 (0.0435, 0) | 0 (0, 0.047) |
| | Youth Neighborhood Safety | 0.222 (0.191, 0.253) | 0.228 (0.2725, 0.186) | 0.224 (0.188, 0.278) | 0.226 (0.2585, 0.198) | 0.232 (0.202, 0.268) |
| | Youth Positive School Environment | 0.548 (0.5215, 0.572) | 0.535 (0.5705, 0.501) | 0.533 (0.4985, 0.573) | 0.534 (0.565, 0.499) | 0.546 (0.513, 0.573) |
| | Youth School Engagement | 0.439 (0.4095, 0.47) | 0.408 (0.4545, 0.367) | 0.43 (0.3915, 0.473) | 0.436 (0.471, 0.4005) | 0.436 (0.401, 0.469) |

| | | | | | | |
|---|---|---|---|---|---|---|
| | Food Security | -0.09 (-0.1245, -0.053) | -0.063 (-0.023, -0.1235) | -0.095 (-0.1345, -0.0555) | -0.081 (-0.027, -0.124) | -0.102 (-0.142, -0.065) |
| | Ability to Pay Bills | -0.112 (-0.1425, -0.072) | -0.1 (-0.0645, -0.157) | -0.096 (-0.137, -0.051) | -0.1 (-0.037, -0.136) | -0.119 (-0.157, -0.093) |
| | Housing Security | -0.096 (-0.13, -0.056) | -0.079 (-0.0425, -0.133) | -0.084 (-0.13, -0.0405) | -0.079 (-0.032, -0.1225) | -0.099 (-0.135, -0.062) |
| | Access to Medical/Dental | -0.036 (-0.067, -0.0055) | -0.037 (0.001, -0.0855) | -0.034 (-0.0705, 0) | -0.028 (-0.0025, -0.0625) | -0.054 (-0.09, -0.016) |
| **Latent Factor 3** | Total Prenatal Conditions | -0.428 (-0.4625, -0.391) | -0.428 (-0.369, -0.483) | -0.416 (-0.475, -0.3555) | -0.427 (-0.472, -0.377) | -0.426 (-0.469, -0.379) |
| | Planned Pregnancy | -0.064 (-0.0975, -0.0325) | -0.07 (-0.019, -0.117) | -0.057 (-0.1, -0.007) | -0.052 (-0.0935, -0.0095) | -0.062 (-0.098, -0.023) |
| | Maternal Age at Birth | -0.1 (-0.134, -0.0655) | -0.102 (-0.055, -0.145) | -0.102 (-0.149, -0.0485) | -0.106 (-0.1475, -0.0665) | -0.105 (-0.145, -0.07) |
| | History of Prenatal Substance Use | 0.054 (0.0095, 0.0945) | 0.062 (0.1225, 0.002) | 0.048 (-0.017, 0.114) | 0.059 (0.017, 0.1065) | 0.062 (0.012, 0.104) |
| | Gestational Age | 0.766 (0.744, 0.7855) | 0.756 (0.786, 0.7245) | 0.744 (0.7195, 0.773) | 0.768 (0.741, 0.795) | 0.768 (0.748, 0.787) |
| | Birth Weight | 0.747 (0.721, 0.7685) | 0.739 (0.773, 0.7045) | 0.722 (0.687, 0.758) | 0.744 (0.7175, 0.7705) | 0.746 (0.722, 0.777) |
| | Youth Family Conflict | 0 (0, 0) | 0 (0, 0) | 0 (0, 0) | 0 (0, 0) | 0 (0, 0) |
| | History of Traumatic Event | 0 (0, 0) | 0 (0, 0) | 0 (0, 0) | 0 (0, 0) | 0 (0, 0) |
| | Parental Poor Adaptive Functioning | 0 (0, 0) | 0 (0, 0) | 0 (0, 0) | 0 (0, 0) | 0 (0, 0) |
| | Highest Parent Education | 0 (0, 0) | 0 (0, 0) | 0 (0, 0) | 0 (0, 0) | 0 (0, 0) |
| | Youth Total Caregiver Acceptance | 0 (0, 0) | 0 (0, 0) | 0 (0, 0) | 0 (0, 0) | 0 (0, 0) |
| | Youth Parental Monitoring | 0 (0, 0) | 0 (0, 0) | 0 (0, 0) | 0 (0, 0) | 0 (0, 0) |
| | Duo Parent Households | 0 (0, 0) | 0 (0, 0) | 0 (0, 0) | 0 (0, 0) | 0 (0, 0) |
| | Sufficient Sleep | 0 (-0.023, 0.0245) | -0.014 (0.02, -0.0555) | 0.007 (-0.018, 0.0445) | 0.005 (-0.0225, 0.0305) | 0.006 (-0.018, 0.03) |
| | BMIz | 0.08 (0.0545, 0.1055) | 0.084 (0.146, 0) | 0.066 (0, 0.1075) | 0.086 (0.0585, 0.117) | 0.076 (0.047, 0.105) |
| | Youth Neighborhood Safety | 0 (-0.026, 0) | 0 (0, 0) | 0 (-0.047, 0) | 0 (-0.0415, 0) | 0 (-0.044, 0) |
| | Youth Positive School Environment | 0 (-0.011, 0) | 0 (0, 0) | 0 (-0.0305, 0) | 0 (-0.0235, 0) | 0 (-0.03, 0) |
| | Youth School Engagement | 0 (-0.027, 0) | 0 (0, 0) | 0 (-0.066, 0) | 0 (-0.0285, 0) | 0 (-0.041, 0) |

| | | | | | | |
|---|---|---|---|---|---|---|
| | Food Security | 0 (0, 0) | 0 (0, 0) | 0 (0, 0) | 0 (0, 0) | 0 (0, 0) |
| | Ability to Pay Bills | 0 (0, 0) | 0 (0, 0) | 0 (0, 0) | 0 (0, 0) | 0 (0, 0) |
| | Housing Security | 0 (0, 0) | 0 (0, 0) | 0 (0, 0) | 0 (0, 0) | 0 (0, 0) |
| | Access to Medical/Dental | 0 (0, 0) | 0 (0, 0) | 0 (0, 0) | 0 (0, 0) | 0 (0, 0) |

**Supplementary Table 4.** Results of models predicting total cortical surface area. A log-likelihood test compared the change in $R^2$ between each model and the null model (fixed + random effects only).

| Total Cortical Surface Area | | | | | |
|---|---|---|---|---|---|
| | **Model 1: Income-to-needs** | **Model 2: Latent Factor 1 + Income-to-needs** | **Model 3: Latent Factor 2 + Income-to-needs** | **Model 4: Latent Factor 3 + Income-to-needs** | **Model 5: Latent Factors 1, 2, & 3 + Income-to-needs** |
| $R^2$ | 0.276 | 0.288 | 0.29 | 0.289 | 0.303 |
| $\Delta R^2$ | 0.0117 | 0.014 | 0.0124 | 0.027 | 0.030 |
| Chi-square | 120.66 | 162.68 | 130.46 | 236.03 | 282.52 |
| *p-value* | < 0.001 | < 0.001 | < 0.001 | <0.001 | <0.001 |
| ***s*(income-to-needs)** | | | | | |
| *F (edf)* | 34.82 (3.77) | 13.17 (3.07) | 34.73 (3.81) | 35.22 (3.71) | 13.91 (3.10) |
| *p-value* | < 0.001 | < 0.001 | < 0.001 | < 0.001 | < 0.001 |
| | | | **Standardized betas (95% CI)** | | |
| Latent Factor 1 | -- | 0.086 (0.06, 0.112) | -- | -- | 0.081 (0.055, 0.107) |
| Latent Factor 2 | -- | -- | 0.033 (0.012, 0.053) | -- | 0.027 (0.007, 0.047) |
| Latent Factor 3 | -- | -- | -- | 0.123 (0.101, 0.145) | 0.121 (0.099, 0.143) |
| Age | 0.003 (-0.014, 0.02) | 0.004 (-0.012, 0.021) | 0.001 (-0.016, 0.018) | 0.004 (-0.012, 0.021) | 0.005 (-0.012, 0.021) |
| Sex | -0.905 (-0.94, -0.87) | -0.909 (-0.943, -0.874) | -0.914 (-0.949, -0.879) | -0.898 (-0.932, -0.863) | -0.908 (-0.943, -0.873) |
| Race-Ethnicity 1[a] | -0.129 (-0.193, -0.066) | -0.126 (-0.189, -0.063) | -0.128 (-0.191, -0.065) | -0.123 (-0.186, -0.06) | -0.12 (-0.182, -0.057) |
| Race-Ethnicity 2[a] | 0.329 (0.266, 0.393) | 0.318 (0.254, 0.381) | 0.332 (0.268, 0.395) | 0.326 (0.263, 0.389) | 0.317 (0.254, 0.38) |
| Race-Ethnicity 3[a] | -0.104 (-0.203, -0.005) | -0.079 (-0.178, 0.019) | -0.107 (-0.206, -0.009) | -0.108 (-0.206, -0.01) | -0.087 (-0.185, 0.011) |
| Race-Ethnicity 4[a] | -0.258 (-0.343, -0.172) | -0.229 (-0.315, -0.143) | -0.259 (-0.345, -0.174) | -0.261 (-0.347, -0.176) | -0.236 (-0.321, -0.15) |

Note: sex was dummy coded as 0 = Male and 1 = Female.
[a]Reference group =Race-Ethnicity5.

**Supplementary Table 5.** Results of models predicting total cognition scores. A log-likelihood test compared the change in $R^2$ between each model and the null model (fixed + random effects only).

| Total Cognition Scores | | | | | |
|---|---|---|---|---|---|
| | **Model 1: Income-to-needs** | **Model 2: Latent Factor 1 + Income-to-needs** | **Model 3: Latent Factor 2 + Income-to-needs** | **Model 4: Latent Factor 3 + Income-to-needs** | **Model 5: Latent Factors 1, 2, & 3 + Income-to-needs** |
| $R^2$ | 0.281 | 0.292 | 0.282 | 0.287 | 0.300 |
| $\Delta R^2$ | 0.064 | 0.076 | 0.066 | 0.071 | 0.083 |
| Chi-square | 557.57 | 672.81 | 576.89 | 597.95 | 726.95 |
| *p-value* | < 0.001 | < 0.001 | < 0.001 | <0.001 | <0.001 |
| ***s*(income-to-needs)** | | | | | |
| *F (edf)* | 94.13 (6.43) | 39.04 (6.17) | 94.51 (6.46) | 95.61 (6.40) | 40.32 (6.18) |
| *p-value* | < 0.001 | < 0.001 | < 0.001 | < 0.001 | < 0.001 |
| | | | **Standardized betas (95% CI)** | | |
| Latent Factor 1 | -- | 0.149 (0.122, 0.176) | -- | -- | 0.145 (0.118, 0.173) |
| Latent Factor 2 | -- | -- | 0.049 (0.027, 0.071) | -- | 0.042 (0.02, 0.063) |
| Latent Factor 3 | -- | -- | -- | 0.075 (0.052, 0.098) | 0.073 (0.051, 0.096) |
| Age | 0.002 (-0.015, 0.019) | 0.299 (0.281, 0.318) | 0.302 (0.28358, 0.32) | 0.301 (0.283, 0.319) | 0.301 (0.283, 0.319) |
| Sex | -0.908 (-0.943, -0.873) | 0.066 (0.029, 0.103) | 0.06 (0.02364, 0.097) | 0.071 (0.034, 0.108) | 0.054 (0.017, 0.091) |
| Race-Ethnicity 1[a] | -0.253 (-0.313, -0.194) | 0.094 (0.029, 0.16) | 0.101 (0.03558, 0.166) | 0.098 (0.033, 0.163) | 0.106 (0.042, 0.171) |
| Race-Ethnicity 2[a] | -0.624 (-0.691, -0.556) | 0.278 (0.212, 0.344) | 0.258 (0.19239, 0.323) | 0.276 (0.211, 0.341) | 0.259 (0.194, 0.324) |
| Race-Ethnicity 3[a] | -0.206 (-0.354, -0.058) | -0.356 (-0.459, -0.254) | -0.314 (-0.41628, -0.212) | -0.36 (-0.462, -0.258) | -0.324 (-0.425, -0.222) |
| Race-Ethnicity 4[a] | -0.237 (-0.302, -0.172) | -0.46 (-0.548, -0.371) | -0.408 (-0.49654, -0.32) | -0.465 (-0.553, -0.377) | -0.418 (-0.505, -0.33) |

Note: sex was dummy coded as 0 = Male and 1 = Female.
[a]Reference group =Race-Ethnicity5.
